# A novel maize gene, *glossy6* involved in epicuticular wax deposition and drought tolerance

**DOI:** 10.1101/378687

**Authors:** LI Li, Yicong Du, Cheng He, Charles R. Dietrich, Jiankun Li, Xiaoli Ma, Rui Wang, Qiang Liu, Sanzhen Liu, Guoying Wang, Patrick S Schnable, Jun Zheng

## Abstract

Epicuticular waxes, long-chain hydrocarbon compounds, form the outermost layer of plant surfaces in most terrestrial plants. The presence of epicuticular waxes protects plants from water loss and other environmental stresses. Cloning and characterization of genes involved in the regulation, biosynthesis, and extracellular transport of epicuticular waxes on to the surface of epidermal cells have revealed the molecular basis of epicuticular wax accumulation. However, intracellular trafficking of synthesized waxes to the plasma membrane for cellular secretion is poorly understood. Here, we characterized a maize glossy (*gl6*) mutant that exhibited decreased epicuticular wax load, increased cuticle permeability, and reduced seedling drought tolerance relative to wild type. We combined an RNA-sequencing based mapping approach (BSR-Seq) and chromosome walking to identify the *gl6* candidate gene, which was confirmed via the analysis of multiple independent mutant alleles. The *gl6* gene represents a novel maize glossy gene containing a conserved, but uncharacterized domain. Functional characterization suggests that the GL6 protein may be involved in the intracellular trafficking of epicuticular waxes, opening a door to elucidating the poorly understood process by which epicuticular wax is transported from its site of biosynthesis to the plasma membrane.

**SIGNIFICANCE STATEMENT:** Plant surface waxes provide an essential protective barrier for terrestrial plants. Understanding the composition and physiological functions of surface waxes, as well as the molecular basis underlying wax accumulation on plant surfaces provides opportunities for the genetic optimization of this protective layer. Genetic studies have identified genes involved in wax biosynthesis, extracellular transport, as well as spatial and temporal regulation of wax accumulation. In this study, a maize mutant, *gl6* was characterized that exhibited reduced wax load on plant surfaces, increased water losses, and reduced seedling drought tolerance compared to wild type controls. The *gl6* gene is a novel gene harboring a conserved domain with an unknown function. Quantification and microscopic observation of wax accumulation as well as subcellular localization of the GL6 protein provided evidence that *gl6* may be involved in the intracellular trafficking of waxes, opening a door for studying this necessary yet poorly understood process for wax loading on plant surfaces.

## INTRODUCTION

The hydrophobic cuticle covers most aerial parts of land plants, which acts as a barrier to protect plants from non-stomatal water loss, ultraviolet (UV) light, physical damage caused by insects or fungi, and from other biotic or abiotic stresses (Shepherd and Wynne Griffiths, 2006). The cuticle mainly consists of two types of lipophilic materials, cutin and cuticular wax. Cutin is the major structural component of the cuticle, which is composed of hydroxy and epoxy C16 and C18 fatty acid monomers, as well as glycerol (Nawrath, 2006). Cuticular waxes, as the second component of the cuticle, are either interspersed in the cutin matrix (cuticular waxes) or overlay the outermost surface of cutin polymer (epicuticular waxes). Cuticular and epicuticular waxes consist of complex mixtures of hydrophobic compounds, mostly very-long-chain fatty acids (VLCFAs) with more than 20 carbon atoms and their derivatives, including primary and secondary alcohols, aldehydes, alkanes, ketones and wax esters (Lemieux, 1996, Kunst and Samuels, 2003, Samuels *et al.*, 2008). The cuticular wax composition and amounts vary greatly among plant species and tissues or organs, as well as development states (Lee and Suh, 2015). Primary alcohols and aldehydes are the major components of the epicuticular waxes in juvenile maize leaves (Javelle *et al.*, 2010).

Epicuticular wax biosynthetic pathways have been extensively studied in *Arabidopsis* by identification and functional characterization of wax-deficient mutant genes (Lee and Suh, 2013). The first step in wax biosynthesis is the elongation of C_16_ and C_18_ fatty acids in the endoplasmic reticulum (ER) into VLCFAs by joining C_2_ building blocks of acetyl-coenzyme A into straight-chain of up to 34 carbon atoms via a fatty acid elongase (FAE) complex (Samuels *et al.*, 2008, Kunst and Samuels, 2009). Following elongation, VLCFAs are modified into different wax products via the distinct alcohol-forming and the -forming pathways (Bernard and Joubes, 2013).

Secretion of epicuticular waxes was elucidated via the identification of two membrane-located ATP binding cassette (ABC) transporters, CER5 and WBC11, responsible for wax export across the PM (Pighin *et al.*, 2004, Bird *et al.*, 2007). However, the mechanism for intracellular trafficking of wax components from their site of synthesis at the ER to the PM is less clear. Through the characterization of the *Arabidopsis ltpg* mutant, lipid transfer proteins (LTPs) have been proposed to be involved in epicuticular wax deposition (Debono *et al*., 2009). In addition, deficient wax secretion in mutants of the *Arabidopsis GNL1* or *ECH* genes that function in endomembrane vesicle trafficking indicated that both genes are involved in intracellular wax trafficking (McFarlane *et al*., 2014).

In maize (*Zea mays*), cuticular wax biosynthesis and accumulation have been aided by the identification of more than 30 glossy (*gl*) loci (Schnable *et al.*, 1994; and unpublished data from the Schnable lab), some of which have been cloned. *gll* and *gl2*, homologs of *Arabidopsis CER3/WAX2* and *CER2*, are involved in leaf epicuticular wax alkane-forming pathway and the extension of VLCFAs to C_30_, respectively (Tacke *et al.*, 1995, Negruk *et al.*,1996, Hansen *et al.*, 1997, Sturaro *et al.*, 2005). *gl4* and *gl8*, homologs of *KSC6* and *KCR* of *Arabidopsis*, belong to the fatty acid elongase (FAE) complex and play important roles in VLCFAs synthesis (Xu *et al.*, 1997, Dietrich *et al.*, 2005, Liu *et al.*, 2009), respectively. *gl3* and *gl15* encode MYB and APETALA2 (AP2)-like transcription factors respectively, and both function in the regulation of epicuticular wax biosynthesis (Moose and Sisco, 1996, Liu *et al.*, 2012). Recently, the *gl13* gene was identified to encode a putative ABC transporter involved in the transport of epicuticular waxes (Li *et al.*, 2013).

One of the most important functions of plant epicuticular waxes is to serve as a protective barrier against environmental stresses, including drought. Drought stress alters the composition and increases the content of epicuticular waxes in *Arabidopsis*, rice and wheat in some cases, and epicuticular wax content has been associated with drought tolerance (Aharoni *et al.*, 2004, Kosma *et al.*, 2009, Zhu and Xiong, 2013, Zhang *et al.*,2015). Some genes involved in biosynthesis and transport of cuticular wax have been proven in improving plant drought tolerance in *Arabidopsis* and several crops (Lee and Suh, 2015, Xue *et al.*, 2017). However, the role of cuticular wax accumulation in drought tolerance in maize remains unclear.

Here, we report the cloning of the maize *glossy6* (*gl6*) gene that is involved in epicuticular wax accumulation and show that the *gl6* mutant, relative to wild type, exhibits reduced epicuticular wax accumulation, as well as increased cuticle permeability and seedling drought sensitivity. Functional characterization indicated that the GL6 protein might be involved in intracellular transport of cuticular wax, providing novel insight into the epicuticular wax biosynthesis/transport pathway.

## RESULTS

### Morphological and biochemical characterization of the *gl6* mutant

The spontaneous *gl6* mutant first described by Emerson in 1935 has been designated *gl6-ref* (Emerson *et al*, 1935, Schnable *et al*, 1994). Like other glossy mutants in maize, seedling leaves of the *gl6-ref* mutant are shiny green in appearance, and water droplets easily form and adher to leaf surfaces after leaves are sprayed with water (Figure 1a). The accumulation of epicuticular waxes on the second leaf surface of *gl6-ref* mutants was examined via Field Emission Scanning Electron Microscopy (FE-SEM). Substantially fewer wax crystals were observed on surfaces of *gl6-ref* mutant leaves relative to surfaces of wild type leaves (Figure 1b). Ultrastructure analysis of leaf epidermal cells by transmission electron microscope (TEM) found acerose inclusions in epidermal cells of *gl6-ref* mutants (Figure 1d, e), but not in wild-type epidermal cells (Figure 1c). This finding is similar to the observation of linear inclusions in *Arabidopsis cer5* mutants which exhibit defects in export of waxes through the PM (Pighin *et al.*, 2004). Our results, therefore, suggested that waxes have accumulated within *gl6* cells.

**Figure 1.**
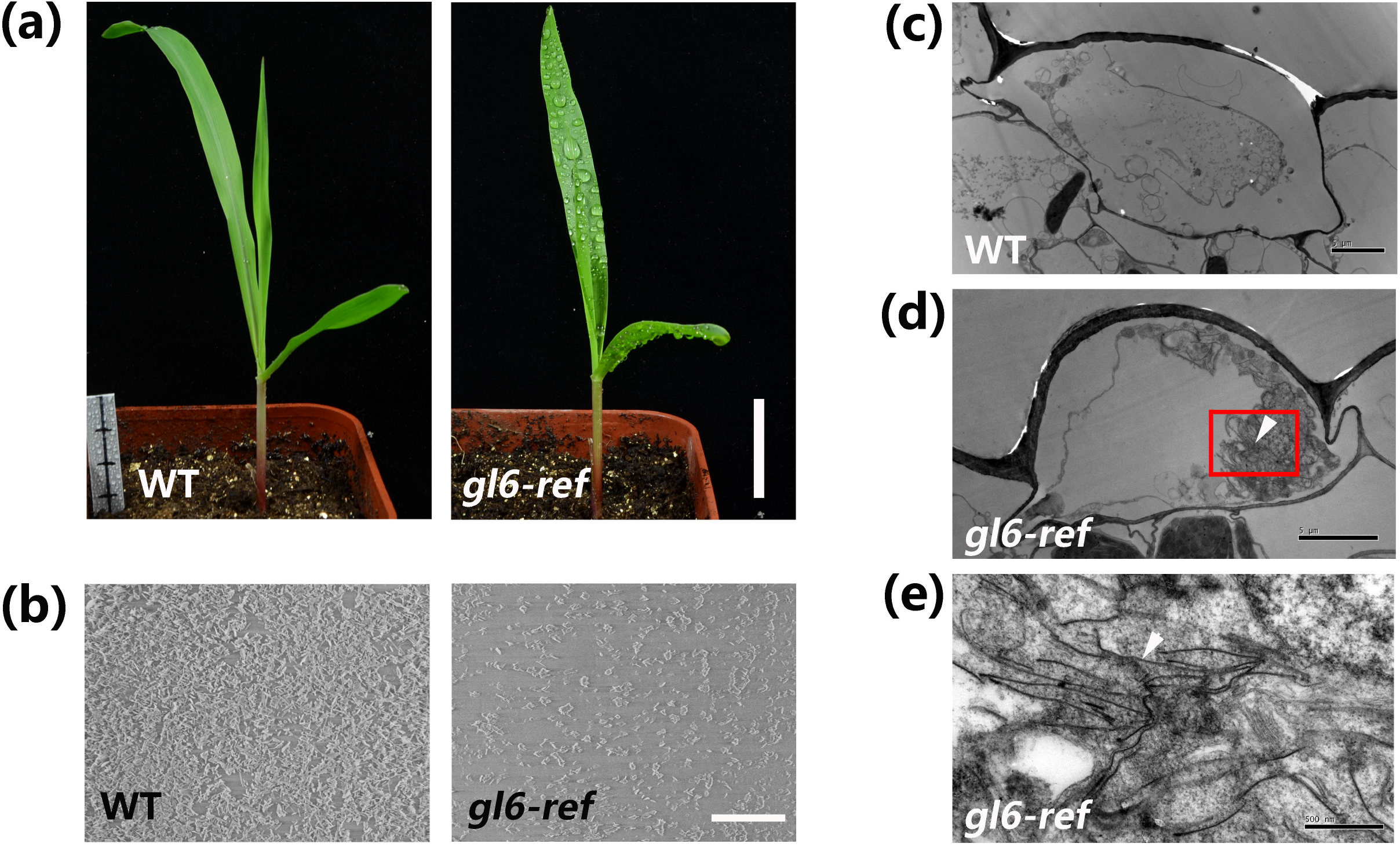
Morphological characterization of the *gl6* mutant. (a) *gl6-ref* mutant exhibits a glossy phenotype. Water is sprayed on seedlings to distinguish *gl6* mutant seedlings from wild type (WT). (b) Leaf epicuticular wax accumulation in WT and mutant (*gl6-ref*) seedlings detected via SEM. 5000×magnification. (c) to (e), TEM analysis of leaf epidermal wax secreting cells wild type (WT) and *gl6-ref* mutant. (e) Enlarged view showing the red rectangle marked area in (d). Arrows in (d) and (e) indicate unusual linear inclusions. Bars = 3 cm in (a), 5 μm in (b), (c) and (d), 500 nm in (e).

The wax load and composition of *gl6-ref* mutants were assessed via GC-mass spectrometry. Relative to wild type, leaf surface waxes loads on *gl6* mutants were reduced by 80%, from 13.2 μg/cm^2^ wild-type level to 2.6 μg/cm^2^ (Figure 2a). Wax composition analysis showed that aldehydes and primary alcohols were highly reduced in *gl6* mutants, which decreased to 1.5% and 22% of wild-type levels, respectively. In addition, the accumulations of fatty acids, alkanes, ethyl stearate, β-sitosterol, and several unidentified wax classes were also decreased in *gl6* mutants (Figure 2b). For individual wax constituents, amounts of C_32_ aldehydes and C_32_ primary alcohols on *gl6* mutant leaves were significantly less than on wild-type leaves. Amounts of C_18_, C_31_ fatty acids, C_29_, C_31_, C_32_ alkanes, C_30_, C_34_ aldehydes and C_30_, C_33_ primary alcohols were also reduced in *gl6* mutants (Figure 2b).

**Figure 2.**
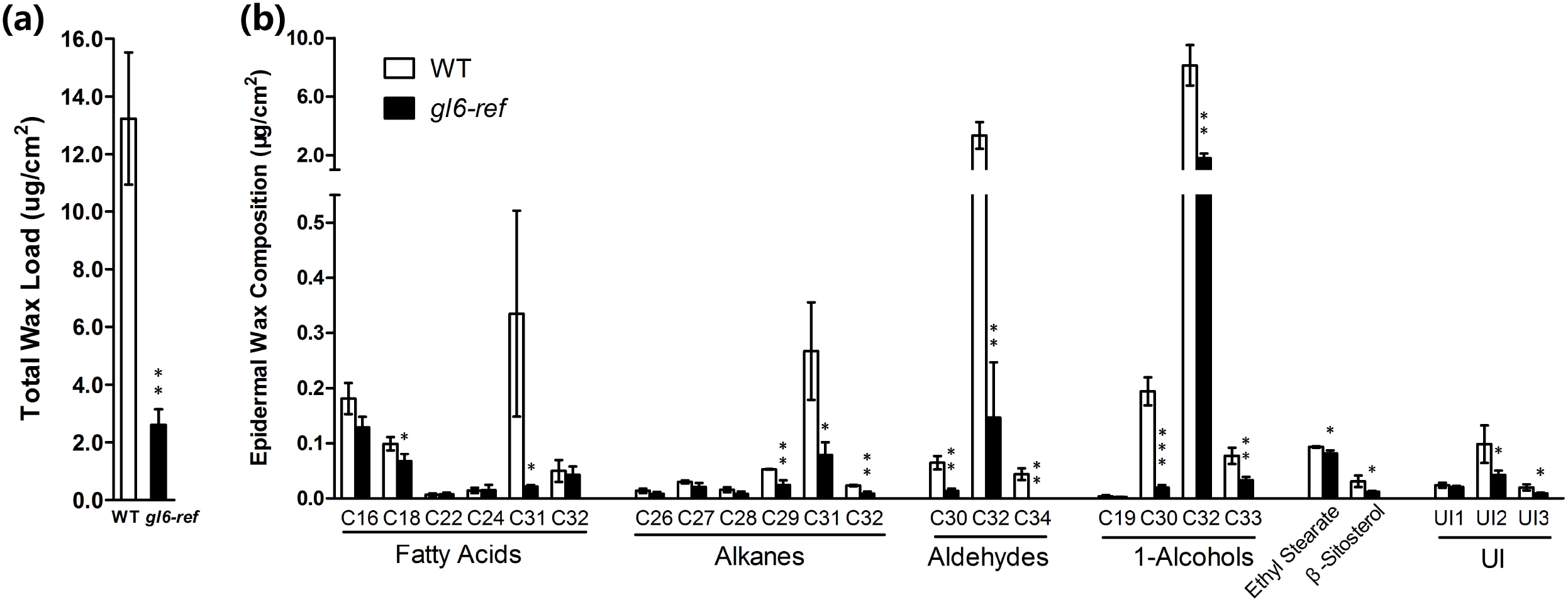
Total leaf epicuticular wax load and wax composition of wild type and *gl6-ref* mutant. (a) Total leaf cuticular wax load of wild type and gl6-refmutant. (b) Epicuticular wax composition of wild type and *gl6-ref* mutant. Values are means of eight biological replicates ± SD. UI indicates unidentified and asterisks indicate statistically significant differences between wild type and *gl6* mutant (*P < 0.05, **P < 0.01, ***P < 0.001, Student’s t test).

We further quantified total waxes including both surface and intracellular waxes and determined the proportion of total waxes secreted to the surface. Total waxes and the proportion of waxes secreted were determined according to published methods (McFarlane *et al.*, 2014). As a result, total waxes were reduced from the wild type levels of 19.7 μg/cm^2^ to 13.1μg/cm^2^ in the *gl6* mutant (Table 1). In wild-type leaves, 67.1% of total waxes were from surface waxes, suggesting that most waxes were secreted (Table 1). In contrast, only 19.9% of total waxes were surface waxes in mutant leaves. In conjunction with the observation of acerose inclusions in *gl6* epidermal cells (Figure 1d, e), these result suggest that the wax exporting system is dysfunctional, causing a large proportion of waxes to remain within cells. Taken together, these results indicated that *gl6* is involved in both the biosynthesis of very-long-chain waxes, particularly C_32_ aldehydes and C_32_ primary alcohols, and, possibly, the wax transport from the ER to the plant surface.

**Table 1.**
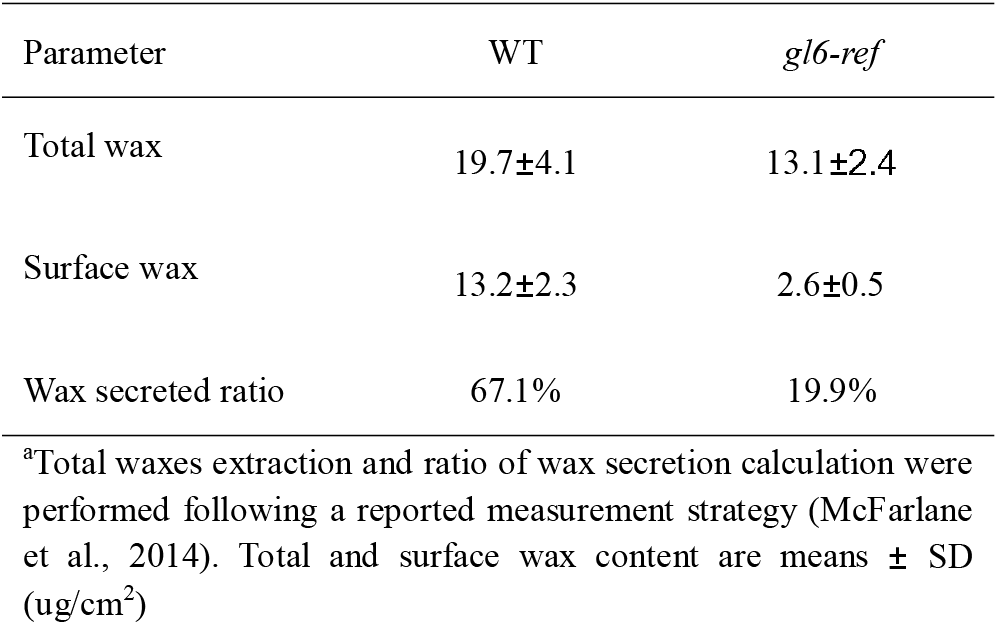
Epicuticular wax extracted from the leaf surface compared with total wax of wild type and *gl6* mutant.^a^

### Increased cuticle permeability and drought sensitivity of *gl6* mutants

Reduced epicuticular wax accumulation is associated with increased cuticle permeability (Kerstiens, 1996, Aharoni *et al.*, 2004, Bessire *et al.*, 2007). We performed chlorophyll leaching and water loss assays using seedling leaves of *gl6-ref* mutants and wild types. For the chlorophyll leaching assay, the concentration of leaf chlorophyll in the solution was monitored at various time points. The results showed that leaf chlorophyll leaching of *gl6-ref* mutants was faster than that of wild type (Figure 3d), even though the total leaf chlorophyll contents of *gl6-ref* mutants and wild types were similar (Figure 3c). To detect water loss, detached leaves of *gl6-ref* and wild type were exposed to air at room temperature; *gl6-ref* mutant leaves were obviously curled and wilted after only 2 hours (Figure 3a). The rate of water loss of detached leaves from *gl6-ref* mutants was significantly higher than that of wild types (Figure 3b). Results from both the chlorophyll leaching and water loss assays suggested that *gl6-ref* mutant leaves exhibit increased cuticle permeability as compared with wild-type leaves.

**Figure 3.**
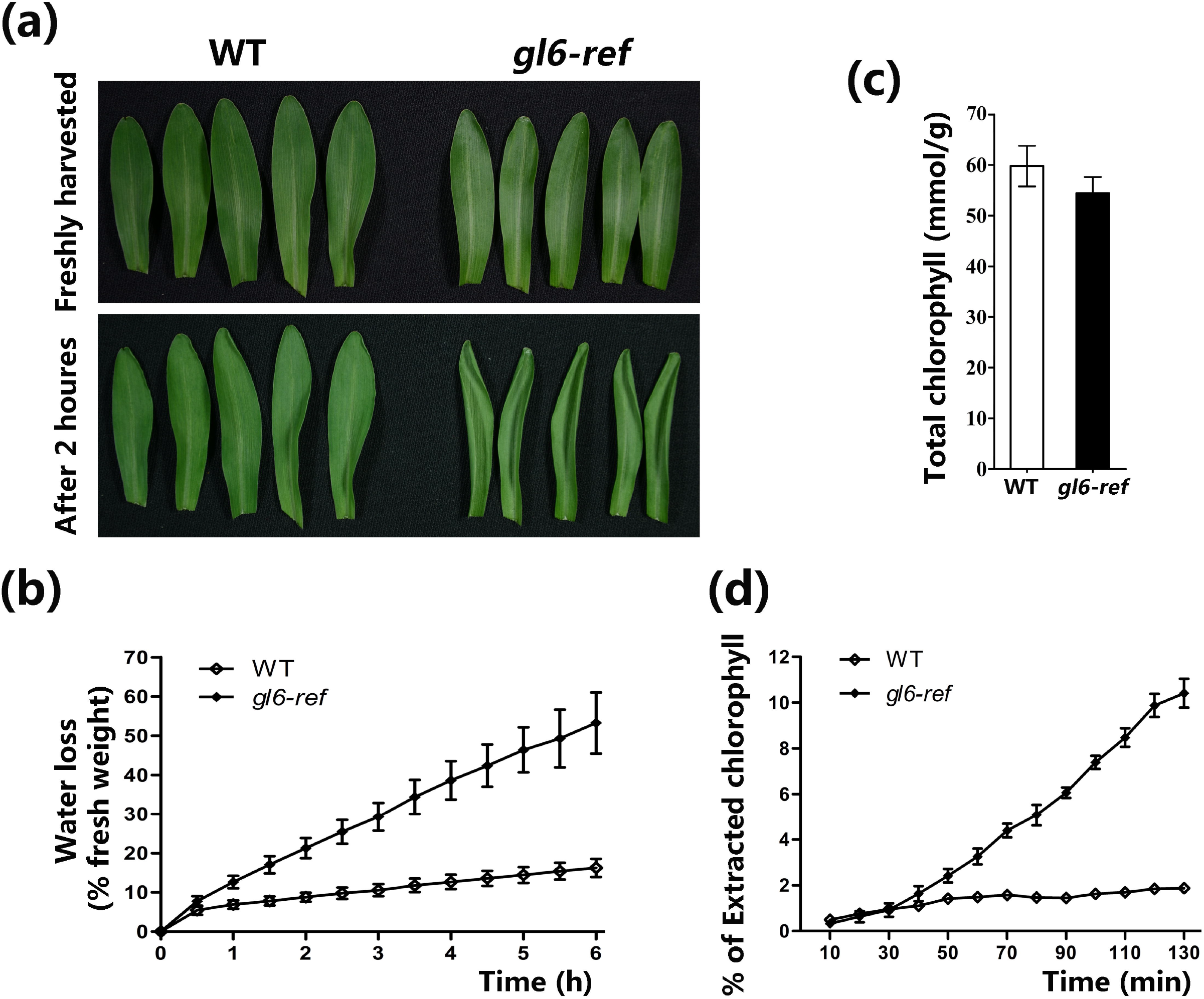
Leaf surface permeability analysis of *gl6* mutant. (a) Detached leaves from WT and *gl6-ref* mutant immediately after harvest and after 2 hours at room temperature. (b) Water loss of detached leaves of the WT and gl6-refmutant. (c) The total leaf chlorophyll content of WT and *gl6-ref* mutant. (d) Kinetics of chlorophyll leaching from leaves of the WT and *gl6-ref* mutant.

Leaf water loss has been associated with reduced leaf surface temperature. Hence, monitoring leaf surface temperature is widely used as an indicator of leaf water loss (Mustilli *et al*., 2002). The surface temperature of *gl6-ref* mutant leaves, monitored via infrared thermography, was lower than that of wild-type leaves under both drought stress and well-watered conditions (Figure 4e), while stomata density, pavement cell density, and stomata index of *gl6-ref* mutants were similar to those of wild type (Figure 4f, g and h). These results suggested that the decreased leaf wax accumulation of *gl6-ref* mutants caused faster water losses.

**Figure 4.**
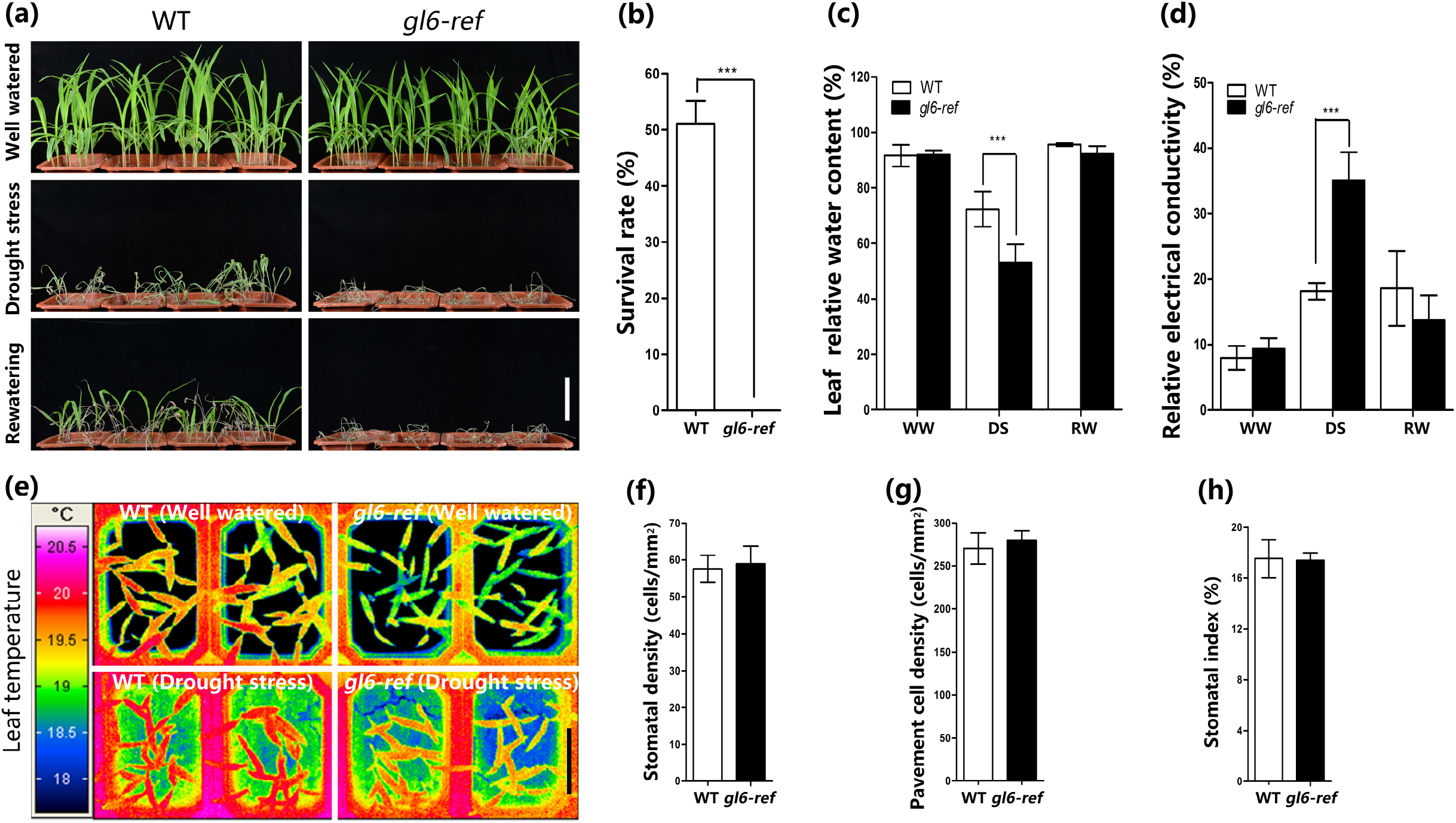
The *gl6* mutant seedlings is sensitive to drought stress. (a) Drought phenotypes of WT and *gl6-ref* mutant seedlings in soil following drought stress and after re-watering. (b) The survival rate of WT and *gl6* mutant seedings after drought stress and re-watering. (c) and (d), The leaf relative water content and relative electrical conductivity of WT and *gl6* mutant seedlings under well watered (WW), drought stressed (DS) and re-watered (RW) conditions. In (b), (c) and (d), data are means of three replicates ± SD (***P < 0.001, Student’s t test). (e) False-color infrared image of the wild type and the *gl6* mutant under well watered and drought stressed conditions. (f) to (h), Stomatal density (f), pavement cell density (g), and stomatal index (number of stomata per total epidermal cells; h) analyzed in leaf abaxial epidermal layers from WT and gl6 mutant. Data are means of five individual plants. Bars = 10 cm in (a) and (e).

Given the increased cuticle permeability and leaf water loss of the *gl6-ref* mutant, we investigated how *gl6-ref* mutant seedlings respond to drought stress. As a result, *gl6-ref* mutants showed a more severe wilting phenotype as compared with wild-type controls (Figure 4a). After re-watering, about 50% of wild-type seedlings could recover and survive, whereas none of *gl6-ref* mutant seedling plants could survive (Figure 4a, b). Under the well-watered conditions, no significant phenotypic differences in either the wilting phenotype or the relative water content of leaves were observed between *gl6-ref* mutants and wild-type seedlings (Figure 4a, 4c). In contrast, under mild-drought stress, the relative water content of *gl6-ref* mutant seedling leaves was lower than that of wild-type controls. The re-watering plants after drought stress showed no significant differences in the leaf relative water content between two genotypes (Figure 4c). In addition, leaf electrolyte leakage assay for evaluating membrane damage was conducted, and the results showed that the *gl6-ref* mutant exhibited a higher relative electrical conductivity than that of wild type under mild-drought stress (Figure 4d), which indicated that a larger degree of membrane damage in mutants resulted in higher electrolyte leakage. In re-watered plants, no significant differences of the relative conductivity were observed between mutants and wild types (Figure 4d). The results corroborated that the *gl6-ref* mutation reduced seedling tolerance to drought.

### Molecular cloning of *gl6*

Previously, the *gl6* gene had been mapped roughly to the long arm of chromosome 3 (Schnable *et al.*, 1994). BSR-Seq, a method utilizing RNA-Seq for bulked segregant analysis (Liu *et al.*, 2012) was performed to map the gene. Briefly, segregating BC_8_F_2_ populations derived from the self-pollination of *gl6-ref* heterozygous plants were used to identify and collect mutant and wild-type plants, separately pooled into two bulks for RNA extraction and sequencing. Three replicates were conducted, producing six RNA-Seq data sets, each of which had more than 20 million reads. RNA-Seq reads were aligned to the B73 reference genome (AGPv2), and the polymorphic SNPs identified which were used as genetic markers to map the target gene. Through this process, the *gl6* gene was mapped to an 18.9 Mb interval on chromosome 3, from 113.5 to 132.4 Mb (Figure 5a).

**Figure 5.**
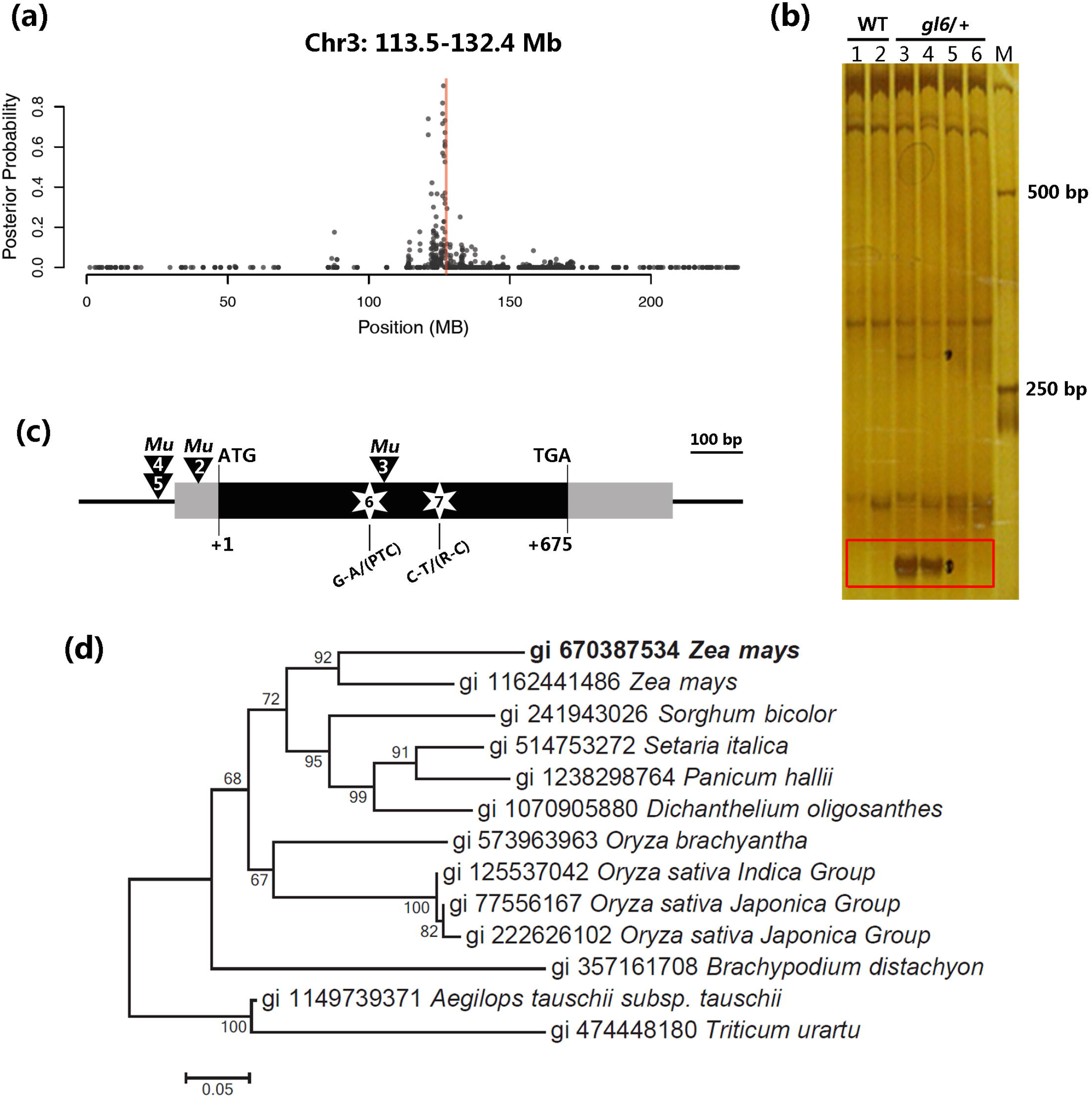
Molecular cloning of the *gl6* gene and its phylogenetic analysis. (a) BSR-Seq analysis of an F_2_ segregated *gl6-ref* population mapped *gl6* to the 113.5-132.4 Mb interval of chr3. (b) Polyacrylamide gel electrophoresis (PAGE) results of Digestion–ligation–amplification (DLA) analysis using the adaptor primer (Nsp-15ctc). The red rectangle indicated the specific bands produced from gl6/+ plants but not in WT plants. (c) The gene structure of *gl6, Mu* insertions in four alleles and lesions in two *gl6* EMS alleles. (d) Phylogenetic tree constructed using MEGA 7.0 and the GL6 protein and GL6 homologs aligned using ClustalW. Distances were estimated with a neighbor-joining algorithm, and bootstrap support is indicated to the left of branches.

To facilitate the identification of the *gl6* gene, additional *gl6* alleles were generated through *Mu* transposon mutagenesis and chemical ethyl methanesulfonate (EMS) treatment (Candela and Hake, 2008). Four new *Mu* tagged alleles were identified in the progeny of crosses between homozygous *gl6-ref* mutants and Mu-active lines; these mutants were designated: *gl6-2, gl6-3, gl6-4* and *gl6-5* (Table S1). Meanwhile, an EMS mutagenesis screen identified three glossy mutants that were verified to be allelic to *gl6-ref*, and which were termed *gl6-6, gl6-7, and gl6-8* (Table S1).

One *Mu* tagged allele, *gl6-2*, was used to perform Seq-Walking sequencing, a genome walking approach to amplify and isolate DNA fragments flanking *Mu* transposon insertions throughout the genome (Li *et al.*, 2013). In total, 28 non-redundant *Mu* flanking sequence (MFS) sites with >30 reads were identified in the 18.9 Mb BSR-Seq mapping interval, and one Mu-insertion hotspot gene, GRMZM2G139786 containing three MFS was identified (Figure S1, Table S2). At the same time, *gl6-2* allele was also subjected to DLA analysis, an adaptor-mediated PCR-based method for isolating *Mu* flanking sequences that co-segregated with mutant phenotypes (Liu *et al.*, 2009). The DLA result independently identified a *Mu* insertion in GRMZM2G139786 that co-segregated the *gl6* glossy phenotype (Figure 5b). Based on these results, all four Mu-derived alleles, *gl6-2, gl6-3, gl6-4* and *gl6-5* were sequenced to identify mutations in GRMZM2G139786. *Mu1* insertions were identified 37 bp upstream of the *gl6* start codon in *gl6-2*, 211 bp upstream of the start codon in both *gl6-4* and *gl6-5* alleles (Dietrich *et al.*, 2002), and 320 bp downstream of the start codon in *gl6-3* allele (Figure 5c).

The genomic sequences of GRMZM2G139786 from three EMS-induced *gl6* alleles were amplified and Sanger sequenced (Table S1). These analyses showed that the *gl6-6* allele contains a G to A transition 314 bp downstream of the start codon, producing a premature termination codon (PTC); *gl6-7* carries a C to T transition 391 bp downstream of the start codon, causing an amino acid change from arginine to cysteine; and the *gl6-8* allele contains a 9-bp deletion 497bp downstream of the start codon and seven non-synonymous mutations in the coding region (Table S1, Figure 5c). In addition, based on the BSR-Seq result, the accumulation of the GRMZM2G139786 transcripts was significantly reduced in the *gl6-ref* mutant pool as compared to the wild-type pool (Table S3). Collectively, these results demonstrate that GRMZM2G139786 is the *gl6* gene.

### Characterization of the novel glossy gene, *gl6*

The *gl6* gene has a single exon with a 675-bp open reading frame and encodes a putative protein with 224 amino acids. Domain analysis showed that GL6 is a novel protein containing a conserved DUF538 (domain of unknown function) between positions 29-138 aa of the protein (Figure S2). Homologs of GL6 can be identified in sorghum, rice, *Arabidopsis* and other plants. Phylogenetic analysis using the full-length protein sequences of GL6 homologs showed that GL6 was grouped into a monocot-specific subfamily (Figure 5d), and multiple sequence alignment showed that the DUF538 domain of these proteins is highly conserved (Figure S2).

A fusion construct of GL6 with C-terminal Yellow Fluorescent Protein (YFP) was generated and expressed in maize protoplast cells. Observation via confocal microscopy showed that GL6-YFP fluorescence signals were was detected in the cytoplasm and the plasma membrane, but not in the nucleus, whereas the control YFP signal was observed throughout the cell (Figures 6a, b). To further study the compartmental localization of the GL6 protein, we co-expressed GL6-YFP with the endoplasmic reticulum (ER) marker red fluorescent protein (RFP)-CNX, the Golgi marker mRFP-ManI, the *trans-Golgi* network (TGN) marker RFP-SYP41 and the nuclear protein marker mRFP-AHL22 in the protoplast cells, and the results revealed that GL6-YFP co-localized with the ER, Golgi and TGN marker (Figures 6c, d and e), but not with the nuclear protein marker (Figure S3). To confirm plasma membrane localization of GL6, the GL6-YFP expressing cells were incubated with the plasma membrane marker FM4-64, and the result showed that the signals of GL6-YFP and FM4-64 were co-localized as expected (Figure 6f). Together, our evidence indicated that GL6 is an ER membrane, Golgi, TGN and plasma membrane-localized protein.

**Figure 6.**
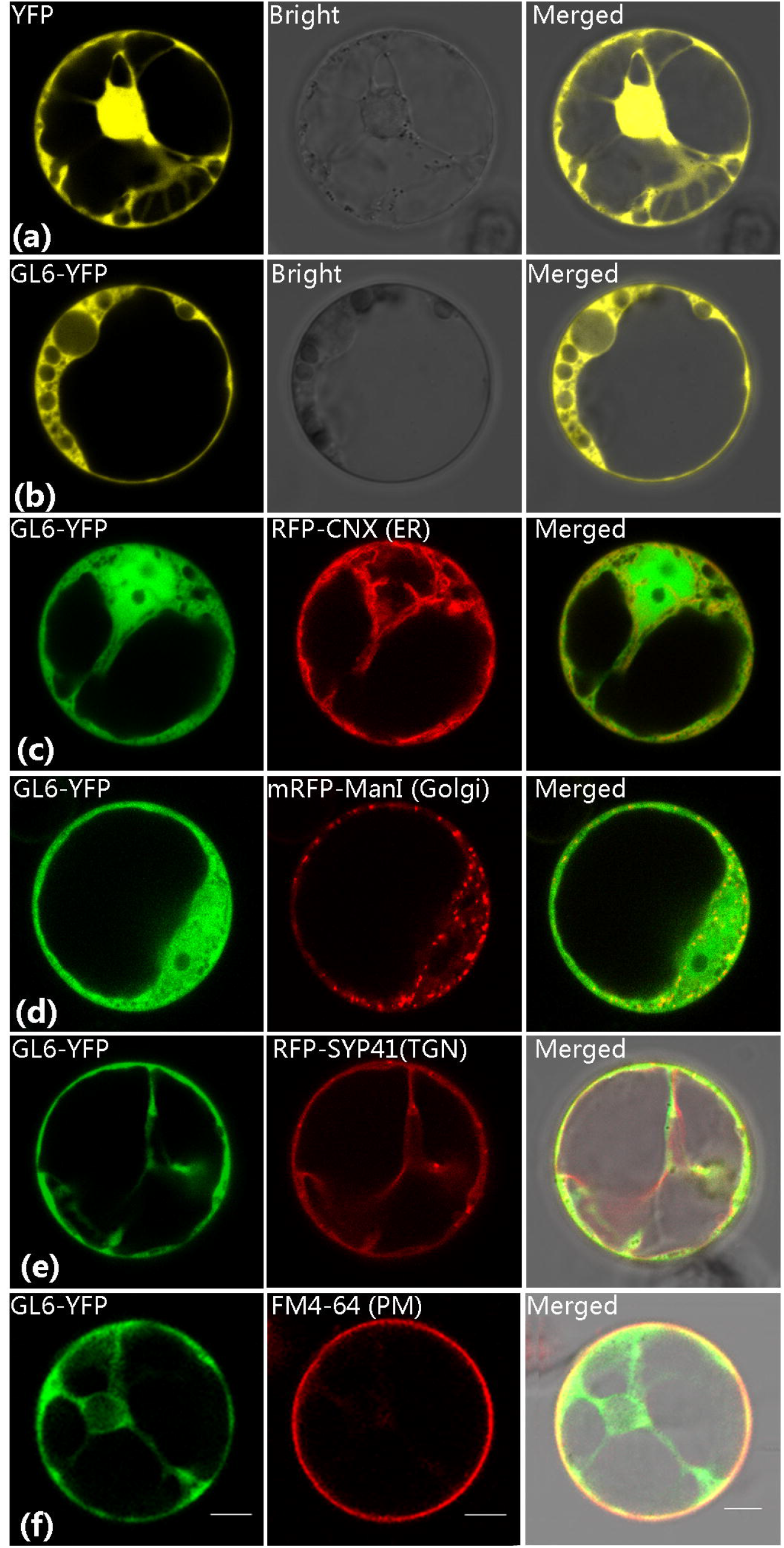
Subcellular localization of the GL6 protein in maize protoplasts. Confocal images show the expression of the YFP, GL6 protein fused at its C terminus with YFP and organelle markers. (a) and (b) Subcellular localization YFP (a) and GL6-YFP (b). (c) Co-localization of GL6-YFP with RFP-CNX (an ER marker). (d) Co-localization of GL6-YFP with mRFP-ManI (a Golgi marker). (e) Co-localization of GL6-YFP with RFP-SYP41(a TGN marker). (f) Co-localization of GL6-YFP with FM4-64(a plasma membrane marker). Bar = 10 μm.

Real-time quantitative RT-PCR analysis was performed to detect *gl6* gene expression in different maize tissues, and the results showed that *gl6* was significantly expressed in leaves and silk, especially in young leaf, but lower expressed in root, husk, anther and immature seed (Figure S4). To further understand the impact of the *gl6* mutation on the transcriptome, we analyzed differential expression in *gl6* mutants and wild types using the BSR-Seq data. With a 10% false discovery rate (FDR), 421 differential expression (DE) genes were identified (Table S4). Among the DE genes, 235 and 186 genes were up- and down-regulated in the *gl6* mutant versus wild type, respectively (Figure 7a). Gene Ontology (GO) enrichment analysis was performed on these DE genes using AgriGO (Du *et al.*, 2010), and complete lists of significantly enriched GO terms are shown in Table S4. Significant enrichment of cellular response to stimulus and stress, especially to water and heat stresses were observed among the enriched GO terms (Figure 7b). Moreover, an enrichment of GO terms related to fatty acid biosynthetic process was also observed. Of genes in these GO terms, two genes, GRMZM2G029912 and GRMZM2G083526, which are homologs of the *Arabidopsis CER3/WAX2* gene (AT5G57800) involved in the biosynthesis of VLCFAs, and another gene, GRMZM2G031790, a homolog of the *Arabidopsis KCS2* gene (AT1G04220), showed significantly higher expression in the *gl6* mutant with 12.0, 4.8 and 12.5 fold changes, respectively (Lee *et al*., 2009, Bernard *et al*., 2012) (Table S3). This result suggested that altering wax amount and localization in the *gl6* mutant resulted in feedback, causing up-regulation of some genes in the fatty acid biosynthesis pathway.

**Figure 7.**
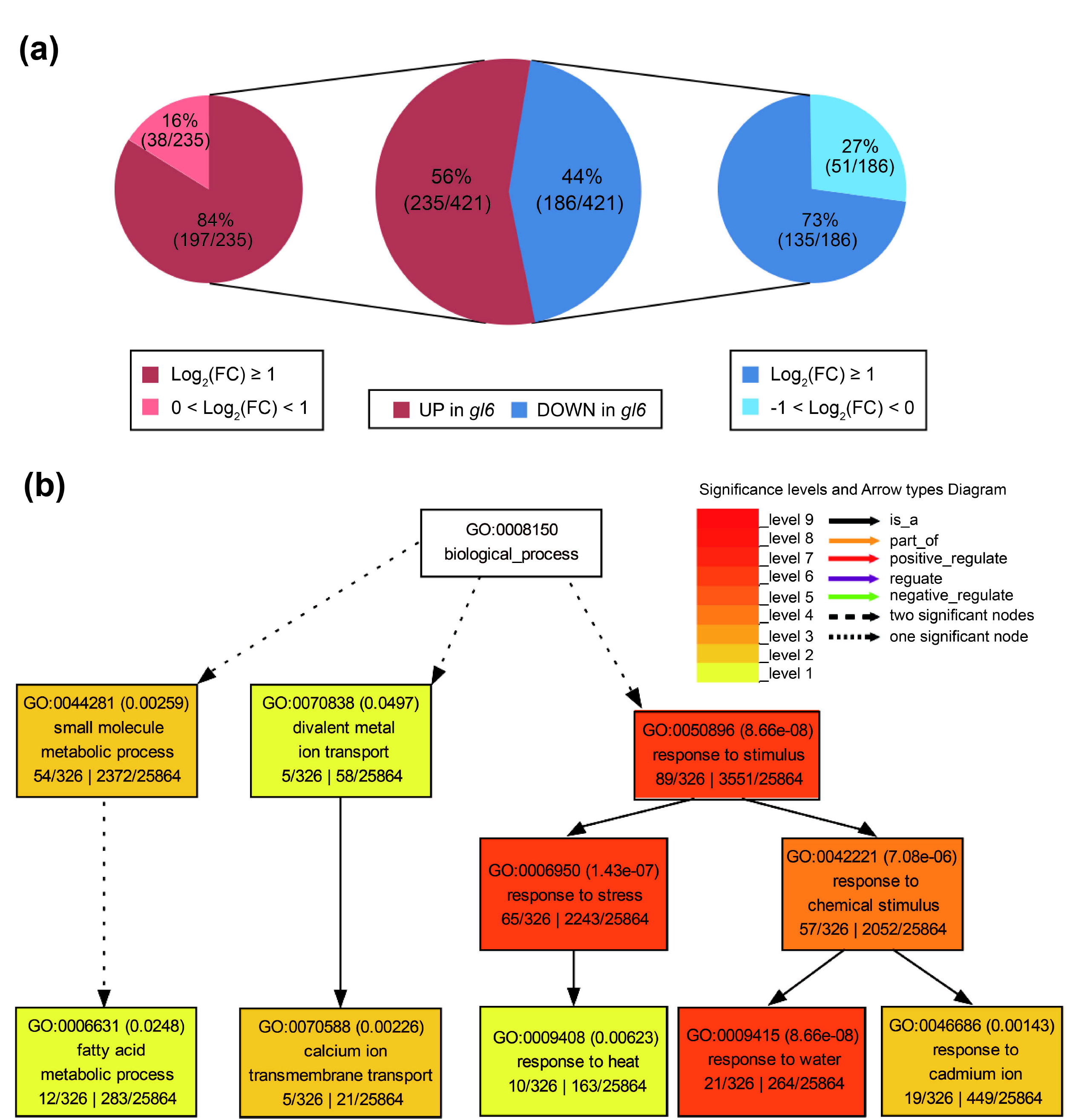
Summary of DE genes and GO Analysis. (a) Summary of the 421 genes that are differentially expressed between *gl6* and WT. The red and blue slices of the pie represent DE genes up- and down-regulated in the *gl6* mutant, respectively. The different shades designate fold changes. (b) GO enrichment analysis of the 421 DE genes. The GO analysis was performed using AgriGO (Du *et al.*, 2010). The processes with deeper color exhibit more significant enrichment. The solid arrows means the process is the only downstream process of the upstream process, while dotted arrows means the process is one of multiple the downstream processes.

## DISCUSSION

In this study, multiple strategies were combined to facilitate the genetic mapping and the identification of a *gl6* candidate gene, which was confirmed using multiple independent *Mu*-tagged and EMS-induced *gl6* alleles. The *gl6* is a novel gene with a conserved uncharacterized domain, opening a door to extending our understanding of the molecular basis of epicuticular wax accumulation.

The phenotype and physiological characterization of the *gl6* mutant showed that *gl6* is a typical maize glossy mutant with reduced epicuticular wax accumulation on the leaf surface, resulting in a glossy phenotype. Epicuticular waxes, the hydrophobic layer on plant surfaces, protect leaves from non-stomatal water loss (Samuels *et al*., 2008). However, few studies provide quantification data to measure drought tolerance of maize glossy mutants. We demonstrated the reduction of epicuticular waxes in the *gl6* mutant was associated with increased cuticle permeability and reduced drought tolerance of maize seedlings. Studies from other plant species have shown that increases in epicuticular wax load can enhance drought tolerance (Lu *et al.*, 2012, Luo *et al.*, 2013, Zhou *et al.*, 2013, Zhu and Xiong, 2013, Al-Abdallat *et al.*, 2014). However, epicuticular wax does not accumulate to high lvels on the leaves of adult maize plants. Future studies should test the impact of elevated expression of epicuticular wax on plant drought tolerance in adult plants as well as the potential physiological cost of accumulating abundant epicuticular waxes.

Sequence analysis revealed that *gl6* encodes a novel protein containing a conserved DUF538 domain with no defined function. The *gl6* orthologs were identified in sorghum, rice, *Arabidopsis* and other plants, but none has been characterized. In this study, we determined that maize *gl6* is involved in wax accumulation and drought tolerance, which will provide insights into understanding the molecular function of DUF538 family members in other plants. In *Arabidopsis*, a wax-deficient phenotype is associated with organ fusion including different abnormalities in trichome development (Wellesen *et al.*, 2001, Chen *et al.*, 2003, Kurata *et al.*, 2003). A T-DNA insertion mutation of the *gl6* homologous gene At1g56580, named *svb-1*, has been examined in *Arabidopsis*,finding smaller trichome with variable branches in *svb-1* mutants versus wild types (Marks *et al.*, 2009). However, no phenotype associated with epicuticular wax accumulation in the *svb-1* mutant was reported, which is worth further investigation on the basis of our findings.

The *gl6* gene encodes a novel protein. Our characterizations have provided clues as to its function. VLCFAs are synthesized in the ER but the processes by which wax components from the ER are delivered to the plasma membrane remain unknown. Two hypotheses have been proposed, Golgi-mediated vesicular trafficking pathway and transfers via physical ER-PM connections (Kunst and Samuels, 2003, Schulz and Frommer, 2004). Two vesicle-trafficking genes, *GNL1* and *ECH*, have been implicated in wax transport from the ER to the PM (McFarlane *et al.*, 2014), which supports the Golgi-mediated vesicular trafficking hypothesis. In the current study, the GL6 protein was localized in the ER membrane, the Golgi, the trans-Golgi network (TGN), and the plasma membrane, which provides further support for the involvement of the Golgi in intracellular wax transport. Our results also suggested that GL6 might play a role in the process of extracellular wax transport. Quantification of secreted waxes on the leaf surface and total leaf waxes showed that a decrease of total waxes in the mutant largely attributed to the reduced accumulation of surface waxes. Ignoring surface waxes, the remaining wax accumulation in mutants was higher than that in wild types, indicative of dysfunctional wax export to leaf surfaces in *gl6* mutants. Acerose inclusions found in epidermal cells of the *gl6* mutant via TEM analysis provided further evidence that waxes accumulate inside epidermal cells of *gl6* mutants. Taken together, our results suggest that GL6 functions in the intracellular transport of cuticular waxes, thereby influencing epicuticular wax accumulation.

## EXPERIMENTAL PROCEDURES

### Plant Materials

The maize (*Zea mays*) *glossy6* reference mutant allele (termed *gl6-ref*, Schnable lab Ac#245) was obtained from in Maize Genetics Stock Center and maintained in the Schnable lab (Schnable *et al.*, 1994). Four additional *glossy6* alleles (*gl6-2, gl6-3, gl6-4 and gl6-5*) isolated via *Mu* transposon tagging screens in Schnable lab from 1992 to 2010 were used to clone the *gl6* gene via Seq-Walking and DLA analysis. Three EMS-induced alleles (*gl6-6* and *gl6-7* were generated by the Schnable lab; *gl6-8* was generated by M.G Neuffer’s lab, GN) were also screened and used to verify the candidate *gl6* gene in this study (Table S1; Schnable *et al.*, 1994).

The *gl6-ref* mutant has been backcrossed to B73 inbred line background up to 8 generations and then continuously self-pollinated. BC_8_F_2_ segregating populations from this backcrossing program were used to map the *gl6* gene by the BSR-Seq technology, and subsequent BC_8_F_3_ homozygous *gl6* mutant or wild type lines were used for the drought tolerance assay, physiology characterization and leaf surface wax analyses.

### Electron Microscopy Techniques

The second leaves collected from *gl6-ref* mutant and wild type were used for FE-SEM analysis according to the standard protocols (Aharoni *et al.*, 2004). In brief, samples were fixed on the spindle and frozen in liquid nitrogen, dried in a vacuum-drying oven and then characterized by FE-SEM microscope (SU8010, HITACHI, Japan).

The collected leaves were also used for TEM analysis by the conventional chemical protocols (Chen *et al*., 2003). Samples were fixed in 2% glutaraldehyde in 0.1 M phosphate buffer (pH 7.4), post-fixed in 1% osmium tetroxide for 1 h, dehydrated with gradient alcohol solutions, and embedded in LR White resin. Ultrathin cross-sections were prepared with a Leica EM UC6 ultramicrotome, stained with 2% uranyl acetate for 2 h, and were analyzed with transmission electron microscope (H-7500, HITACHI, Japan).

To measure the stomatal and pavement cell density and calculate the stomatal index, imprints were detached from the surface of the collected leaves and were mounted on glass microscope (Nadakuduti *et al*., 2012). Samples were observed with an optical microscope (IX71, Olympus, Japan).

### Water Loss and Chlorophyll Leaching Assay

Detached leaves of *gl6-ref* mutant and wild-types were left at room temperature and photographed and weighed every 1 h. Water loss was calculated and represented by the percentage of fresh weight. Chlorophyll leaching assays were carried out according as described (Aharoni *et al.*, 2004). Briefly, excised seedling leaves were washed with tap water, weighed, and put in 30 mL of 80% ethanol at room temperature. Four hundred microliters were taken out from each sample every 10 min, and were used to measure the absorbance at 664 and 647 nm wavelength. Finally, the samples were incubated in boiling water, cooled on ice, and were used to examine the absorbance to evaluate the total chlorophyll content. The formula in the described method was used to calculate the chlorophyll content and chlorophyll extraction rate.

### Drought Tolerance Experiment

Seeds of the BC8F3 homozygous *gl6* mutant or wild type were germinated in paper towels for 2 days and then transplanted in sand-filled pots in a greenhouse (27 °C day/23 °C night, 16 h day). The control plants were well watered, and the treatment plants were subjected to drought stress by by withholding water for up to 20 days, and re-watered. Six biological replications were used for this assay, and the number of surviving seedlings used to calculate the survival rate. During the drought treatment, the relative water content (RWC) and relative electrical conductivity (REC) were respectively monitored by the previously described method (Zheng *et al.*, 2010). Thermal images were obtained using an infrared imaging system (VarioCam HD, InfraTec, Germany) following the manufacturer’s instructions.

### Analysis of Wax Composition

Wax extraction, gas chromatography (GC)-mass spectrometry (MS) analyses were performed according to the described methods with some modifications (Chen *et al.*, 2011). The *gl6* mutants and the wild-types grown in the substrate of roseite and sands (1:1) at growth chamber (25°C) under 16/8 hrs light/dark for three-leaves (about 6-7 days after plant, DAP), the second leaves (6 DAP) contained 5-6 g dry matter from *gl6* mutant and wild type were fresh collected and instantly immersing into 1,000 μl chloroform for 45 s, the extracts containing 10 mg of tetracosane (Fluka) as an internal standard, was transferred into opened reactive vials, dried with nitrogen gas (Pressure Blowing Concentrator; N-EVAP), and derivatized by adding 20 l of N,N-bis-trimethylsilyltrifluoroacetamide (Macherey-Nagel) and 20 l of pyridine and incubated for 40 min at 70°C. These derivatized samples were then analyzed by GC-FID (Agilent, Technologies) and GC-MS (Agilent gas chromatograph coupled to an Agilent 5973N quadrupole mass selective detector).

### BSR-Seq, Seq-Walking and DLA Analysis

About twenty glossy and non-glossy seedling segregants from the *gl6-ref*BC_8_F_2_ population were collected to construct mutant and non-mutant bulks for RNA extraction. Total RNA was isolated using RNeasy Plus Mini Kit (Qiagen, USA) and RNA quality (RIN scores over 8) was checked on a Bioanalyzer 2100 (Agilent technologies, Life technology, USA) using an RNA 6000 Nano chip. RNA-Seq libraries were constructed using the Illumina RNA-Seq sample preparation kit according to the manufacturer’s protocol. The RNA-Seq libraries were sequenced on an Illumina HiSeq2000 instrument. Sequencing data were used to perform BSR-Seq analysis following our published method (Liu *et al.*, 2012). For the Seq-Walking analysis, DNA was extracted using a CTAB method (Murray and Thompson, 1980) from the pools of heterozygous of *B73×(gl6-Mu/gl6-ref)*, in which half of the individuals contains Mu-insertion allele, *gl6-2* (Table S1). Following quantification, DNA samples were sheared using a Biorupter machine (Diagenode, USA) and sequenced. The resulting sequence data were analyzed according to our published Seq-Walking Strategy (Li *et al.*, 2013). DNA samples of *gl6-2* were also used to perform Digestion-Ligation-Amplification (DLA) analysis according to our published method (Liu *et al.*, 2009).

### Transcriptome analysis

RNA-Seq data of the *gl6* mutant and non-mutant bulks used in the BSR-Seq analysis were also used to compare transcriptome-wide differences between the mutant and its wild-type control. Reads were aligned to the maize reference genome B73 AGPv3 using Tophat2 (Kim *et al.*, 2013). Transcript accumulation levels were calculated using the R package HTSeq (Anders *et al.*, 2015) and differentially expressed genes were identified with the R package DESeq2 (Love *et al.*, 2014). Tests for enrichment of differentially expressed genes were performed using agroGO with the standard settings (Fisher’s exact test with significance threshold of 0.05, Yekutieli multi-test adjustment) (Du *et al.*, 2010).

### qRT-PCR

For tissue-specific expression analysis, samples of young roots and leaves of 12 and 24 days after sowing (DAS), mature leaves of 12 and 27 days after pollination (DAP), husk, silk, anther, endosperm, embryo and seeds of 14 and 27 DAP were collected from the inbred line B73, and then were used to extract total RNAs by TRIzol reagent (Invitrogen). Quantitative real-time RT-PCR (qRT-PCR) analysis was conducted with TransStart Green qPCR SuperMix (TransGen Biotech) on a 7300-sequence detection system (Applied Biosystems). Maize *actin1* was used as internal control, and the relative expression of *gl6* mRNA was calculated using the 2^−ΔΔCt^ method (Livak and Schmittgen, 2001).

### Subcellular Location of GL6

To generate the GL6-YFP fusion construct, the open reading frame (ORF) of the *Gl6* gene was cloned into the pEarleyGate-101 vector which contains the yellow fluorescent protein (YFP) reporter gene (Earley *et al.*, 2006). Using previously described PEG-mediated transformation protocols (Chen *et al*., 2014), the GL6-YFP construct was transformed into maize protoplasts alone or co-transformed with a ER marker (red fluorescent protein (RFP)-CNX), a Golgi marker (mRFP-ManI), a trans-Golgi network (TGN) marker (RFP-SYP41), or and a nuclear protein marker (mRFP-AHL22) (Xiao *et al.*, 2009, Cui *et al.*, 2014). Fluorescence signals were observed under a confocal laser scanning system (Leica Microsystems, Wetzlar, Germany).

## ACKNOWLEDGEMENTS

This work was supported by the National Key Research and Development Program of China (Grant No. 2016YFD0101002), the 948 project of the Ministry of Agriculture of China (Grant No. 2015-Z11), the Agricultural Science and Technology Innovation Program of CAAS and the National Natural Science Foundation of China (31701437). We thank Mr. Cheng-Ting “Eddy” Yeh, Drs. Wei Wu, Heng-Cheng Hu and An-Ping Hsia for technical support and helpful discussions, Ms. Mitzi Wilkening for sequencing services in ISU Genomic Technologies Facility, former members of the Schnable lab including the late Joel Hansen and Philip Stinard, and the Schnable Lab’s current nursery manager, Ms. Lisa Coffey for the generation and maintenance of genetic stocks used in this study. We thank Dr. Jianxin Shi, Dr. Guorun Qu, and Ms. Qian Luo from Shanghai Jiao Tong University for their assistance in conducting the GC-FID and GC-MS measurements and associated data analyses.

## SUPPORTING INFORMATION

Figure S1. Mu insertion sites from Seq-walking.

Figure S2. Multiple sequence alignment of amino acid sequences of the predicted maize GL6 protein and its homologs in other species.

Figure S3. Co-localization of GL6-YFP with mRFP-AHL22 (nuclear marker).

Figure S4. Accumulation of gl6 transcripts in various tissues.

Table S1. *gl6* alleles information and the corresponding mutation types

Table S2. *Mu* flanking sequences and *Mu* sequences for each *gl6 Mu* tagging allele

Table S3. Differential expressed genes (DEGs) in gl6 mutant vs. wild-types

Table S4. Enriched GO terms in *gl6* mutant differentially expressed genes

---

Figure S1. *Mu* insertion sites from Seq-walking. Number of reads at each *Mu* insertion site is plotted versus site’s physical coordinates. Red dots highlight *Mu* insertion sites within the *gl6* candidate gene. The region shown covers the mapping interval on chromosome 3.

Figure S2. Multiple sequence alignment of amino acid sequences of the predicted maize GL6 protein and its homologs in other species. Sequences were obtained from NCBI and aligned using the website (http://multalin.toulouse.inra.fr/multalin/multalin.html) following default parameters. Zm, *Zea mays*; Do, *Dichanthelium oligosanthes*; Si, *Setaria italic*; Ph, *Panicum hallii*; Sb, *Sorghum bicolor*; At, *Aegilops tauschii;* Bd, *Brachypodium distachyon*; Ob, *Oryza brachyantha*; Os, *Oryza sativa;* Tu, *Triticum urartu*.

Figure S3. Co-localization of GL6-YFP with mRFP-AHL22 (nuclear marker). Confocal images show the expression of GL6 protein fused at its C terminus with YFP and the nuclear marker mRFP-AHL22. Bar = 10 μm.

Figure S4. Accumulation of *gl6* transcripts in various tissues. Real-time quantitative RT-PCR analysis of roots, leaves, husk, silk, anther and seeds from the inbred line B73, normalized using the maize tublin gene. Values are means of three replicates ± SD. DAS indicates days after sowing, DAP indicates days after pollination.

